# DHHC20 palmitoyl-transferase reshapes the membrane to foster catalysis

**DOI:** 10.1101/808469

**Authors:** R. Stix, J. Song, A. Banerjee, J.D. Faraldo-Gómez

## Abstract

Cysteine palmitoylation, a form of S-acylation, is a key post-translational modification in cellular signaling. This type of reversible lipidation is catalyzed by a family of integral membrane proteins known as DHHC acyltransferases. The first step in the S-acylation process is the recognition of free acyl-CoA from the lipid bilayer. The DHHC enzyme then becomes auto-acylated, at a site defined by a conserved Asp-His-His-Cys motif. This reaction entails ionization of the catalytic Cys. Intriguingly, in known DHHC structures this catalytic Cys appears to be exposed to the hydrophobic interior of the lipid membrane, which would be highly unfavorable for a negatively charged nucleophile, thus hindering auto-acylation. Here, we use biochemical and computational methods to reconcile these seemingly contradicting facts. First, we experimentally demonstrate that human DHHC20 is active when reconstituted in POPC nanodiscs. Microsecond-long all-atom molecular dynamics simulations are then calculated for hDHHC20 and for different acyl-CoA forms, also in POPC. Strikingly, we observe that hDHHC20 induces a drastic deformation in the membrane, particularly on the cytoplasmic side where auto-acylation occurs. As a result, the catalytic Cys becomes hydrated and optimally positioned to encounter the cleavage site in acyl-CoA. In summary, we hypothesize that DHHC enzymes locally reshape the membrane to foster a morphology that is specifically adapted for acyl-CoA recognition and auto-acylation.

**Significance Statement:** Palmitoylation, the most common form of S-acylation and the only reversible type of protein lipidation, is ubiquitous in eukaryotic cells. In humans, for example, it has been estimated that as much as ∼10% of the proteome becomes palmitoylated, i.e. thousands of proteins. Accordingly, protein palmitoylation touches every important aspect of human physiology, both in health and disease. Despite its biological and biomedical importance, little is known about the molecular mechanism of the enzymes that catalyze this post-translational modification, known as DHHC acyltransferases. Here, we leverage the recently-determined atomic-resolution structure of human DHHC20 to gain novel insights into the mechanism of this class of enzymes, using both experimental and computational approaches.

## Introduction

DHHC acyltransferases are integral membrane proteins that catalyze cysteine-acylation (S-acylation), one of the most common post-translational modifications in eukaryotic cells (1, 2). (The term DHHC reflects a consensus Asp-His-His-Cys catalytic motif). Indeed, systems-level analyses estimate that as much as ∼10% of the human proteome becomes palmitoylated, including over 40% of all known synaptic proteins (3). Accordingly, palmitoylation touches every important aspect of human cellular physiology, both in health and disease. Human DHHC proteins have been speficically implicated in the development of multiple cancer types as well as Huntington’s disease, schizophrenia and X-linked mental retardation (2, 4).

S-acylation is unique in that it is the only form of lipidation that can be biochemically reversed, by a class of enzymes known as thioesterases (5, 6). Reversible post-translational modifications are used by the cell to regulate the activity and co-localization (or segregation) of proteins; S-acylation is thus specifically important in membrane signaling (2). S-acylation proceeds in two stages (7). First, a DHHC enzyme embedded in the lipid bilayer binds fatty acyl-CoA, and the active-site cysteine becomes auto-acylated. This process involves the formation of a thioester bond between the fatty acyl chain (hereafter referred to as “acyl chain”) and the cysteine; CoA is cleaved off as a result. In the second stage, the acyl chain covalently linked to DHHC is transferred to a protein substrate (2).

The only human DHHC protein of known structure is DHHC20 (hDHHC20) (8). hDHHC20 is known to be selective for C16:0 acyl chains (8), i.e. it specializes in palmitoylation, which is the most prevalent form of S-acylation (9). Acylation occurs with other chain lengths, but is suboptimal (8). hDHHC20 consists of four transmembrane helices and a cytoplasmic domain (**Fig. 1A**). The available structure (8) does not fully clarify the mode of acyl-CoA recognition; however it does reveal the location of the catalytic Asp-His-His-Cys motif, and what appear to be the binding surface for the acyl chain, in the transmembrane span. Based on these insights, Rana et al. have proposed a catalytic mechanism whereby His154, polarized by Asp153, acts as a base and causes the deprotonation of Cys156. Ionized Cys156 then acts as a thiolate nucleophile and attacks the acyl-CoA carbonyl to create a thioester bond (**Fig. 1B**) (10).

**Figure 1.**
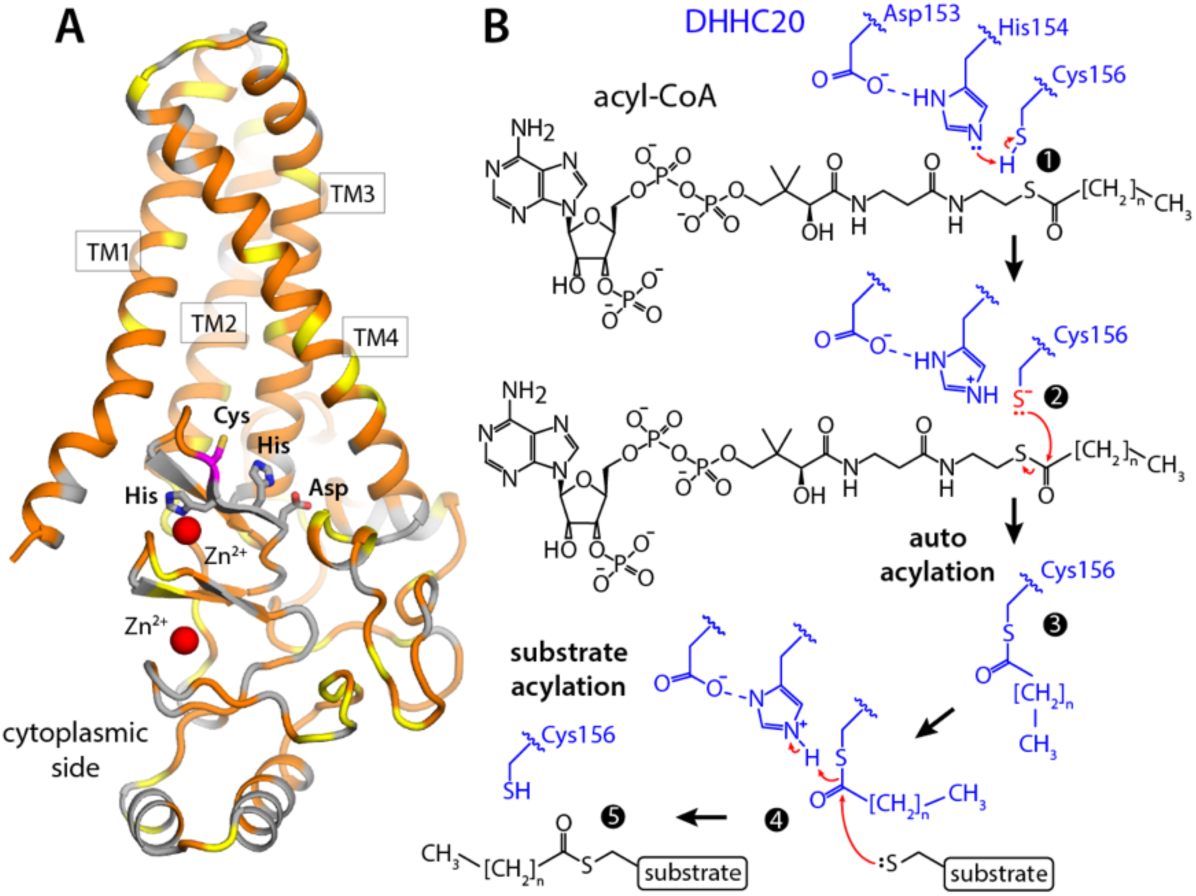
Structure of human DHHC20 and proposed catalytic mechanism. **(A)** Crystal structure of hDHHC20 (PDB entry 6BMN). The overall fold of the structure is represented with cartoons, indicating the 4 transmembrane spans (TM1-TM4). Hydrophobic and/or aromatic residues are colored in orange; serine and threonine residues in yellow; others in gray. The sidechains that the define the catalytic site (Asp-His-His-Cys) are highlighted, as well as two structural Zn^2+^ ions. **(B)** Scheme for the 2-stage S-acylation reaction mediated by DHHC20: steps 1-3 comprise the auto-acylation stage, where by an acyl-chain is linked to Cys156; in steps 4-5, this acyl-chain is transferred to a substrate. Red arrows indicate the proposed reaction pathway.

It has been puzzling, however, that the catalytic Cys appears to be well within the hydrophobic, transmembrane region of the protein. Shielded from water, the proposed deprotonation of this side-chain would be energetically costly, and hinder the auto-acylation reaction. An equally important question is whether this side chain is accessible to the reactive thioester group in free acyl-CoA. Here, we seek to resolve these questions by combining experimental and computational approaches. First, we use biochemical assays to evaluate the functionality of purified hDHHC20 reconstituted in a simple phospholipid bilayer. Using the same type of lipid bilayer, atomically-detailed molecular dynamics simulations are then calculated, both for hDHHC20 and for 3 acyl-CoA variants, to gain insights into this recognition process as it pertains to the membrane.

## Materials and Methods

### Protein expression and purification

Expression and purification of human DHHC20 (hDHHC20) from *Pichia* were carried out as described previously (8), with the following modifications: the hDHHC20 expression construct was cloned into a modified pPICZ-N vector with a His_10_ tag followed by a mVenus coding sequence, and finally, by a PreScission cleavage site at the N-terminus of the hDHHC20-encoding DNA sequence. hDHHC20 was purified with an intact mVenus tag and quantified using nanodrop (ε_max_ = 105,000 M^-1^). MSPE3D1 was produced as described previously (11) with the following modifications: the TEV cleavage site in the original pET28a MSPE3D1 construct (addgene #20066) was mutated to a PreScission cleavage site. The N-terminal His_6_ tag of MSPE3D1 was removed using PreScission.

### Reconstitution in POPC nanodiscs

The hDHHC20 nanodisc reconstitution was carried out in the following condition: 6 µM hDHHC20 mVenus, 18 µM MSPE3D1, 1.44 mM POPC, 20 mM sodium cholate in 25 mM Tris, 137 mM NaCl, 27 mM KCl, pH 7.4 buffer. The reaction mixture was left agitating gently at 4 °C for 2 h, and freshly washed amberlite beads (∼0.8-1 g per reaction volume) were added to capture the residual detergent. The reaction was left agitating gently overnight at 4 °C. The supernatant was filtered through 0.45 um PVDF filter prior to gel filtration on Sepharose 200 Increase column using 25 mM Tris, 137 mM NaCl, 27 mM KCl, pH 7.4 buffer. The fractions containing hDHHC20 nanodiscs were collected and concentrated using milipore 50 kDa concentrators and quantified (ε_max_ = 105,000 M^-1^).

### Palmitoyl-transferase assay in nanodiscs

The palmitoyl-transferase (PAT) assay has been described in detail elsewhere (12). In brief, reaction mixture **1** containing 50 mM HEPES, 50 mM NaCl, 2 mM oxo-ketoglutarate, 0.5 mM NAD^+^ and 0.4 mM TPP was made in 4 °C, with and without hDHHC20 nanodiscs. Mixture **2** was prepared containing 40 µM palmitoyl-CoA, 50 mM HEPES, 50 mM NaCl, and 0.2 mUnits of α-ketoglutarate dehydrogenase, and stored in 4 °C. In 96-well CORNING black-well solid bottom plates (catalog #: 12-566-09), 30 µL of reaction mixture **1** with and without hDHHC20 nanodiscs were plated, DTT was added (varying concentrations from 50 mM to 0.01 mM), and NADH fluorescence kinetics were measured after adding 30 µL of mixture **2** containting palmitoyl-CoA, using TECAN plate reader. The fluorescence of the wells containing hDHHC20 nanodiscs was normalized in reference to that of the wells lacking the enzyme, and the normalized fluorescence was converted to [NADH, µM] using the following equation derived from an NADH calibration curve: RFU = 88.4 × [NADH, µM] + 22.

### 2-BP inhibition assay in nanodiscs

2-bromopalmitate (2-BP) inhibition of hDHHC20 nanodiscs was monitored by varying concentrations of 2-BP in reaction mixture **1** containing hDHHC20 nanodiscs for final concentration range of 65.7 µM to 2.0 nM. DMSO vehicle controls of reaction mixture **1**, both with and without hDHHC20 nanodiscs, were used to normalize the inhibitor treated wells, and the resulting data was plotted and analyzed with GraphPad (non-linear regression, log(inhibitor) v. response - variable slope).

### Molecular dynamics simulations

The simulations of hDHHC20 were based on the available apo-state crystal structure (PDB entry 6BMN) (8). The protein structure was embedded in a pre-equilibrated POPC bilayer using GRIFFIN (13). The simulation system consists of 222 POPC lipids, 21633 water molecules, 31 Na^+^ and 43 Cl^-^ (100 mM NaCl plus counter-ions of the protein net charge), in an orthorhombic box of ∼89 × 89 × 119 Å. The preparation of the simulations entailed a multi-stage equilibration phase whereby a series of (primarily) internal-coordinate restraints acting on different portions of the protein structure are gradually weakened and ultimately removed over 100 ns. A trajectory of 1 µs was then calculated for each of the two equilibrated configurations, with no conformational restraints.

Simulations were also carried of caprylyl-CoA (C8:0), palmitoyl-CoA (C16:0), and behenyl-CoA (C22:0), in a POPC bilayer. These simulation systems consist of 240 POPC lipids, 17887 water (14)molecules, 34 Na^+^ and 30 Cl^-^ (100 mM NaCl plus counter-ions of the acyl-CoA net charge), in an orthorhombic box of ∼87 × 87 × 109 Å. The palmitoyl-CoA simulation was prepared by placing the lipid outside the membrane, in the water layer. Palmitoyl-CoA inserted spontaneously in the bilayer within ∼200 ns, and remained therein for the duration of trajectory, which was 1-µs long. The simulations of caprylyl-CoA and behenyl-CoA were prepared by elongating or shortening the acyl chain in palmitoyl-CoA, once inserted in the membrane.

All MD simulations were carried out using NAMD version 2.9 (15) and the CHARMM36 force-field (16-18), at constant temperature (298 K) and constant pressure (1 atm), with periodic boundary conditions. Electrostatic interactions were calculated using PME, with a real-space cut-off of 12 Å. A cut-off of 12 Å was also used for computing van-der-Waals interactions, with a smooth switching function taking effect at 10 Å. The integration time-step was 2 fs.

Force-field parameters for palmitoyl-CoA (C16:0), caprylyl-CoA (C8:0) and behenyl-CoA (C22:0) were derived from existing CHARMM27 parameters for acetyl-CoA (14). Adaptations were made by incorporating the relevant CHARMM36 parameters for protein and ADP into the acyl-CoA parameter set. All three force-fields are available upon request to the authors.

## Results

### hDHHC20 is active in POPC nanodiscs

We first sought to identify a type of lipid bilayer that is biologically representative and yet simple enough to permit an unequivocal interpretation of molecular simulation data. Given that PC lipids are the most abundant in eukaryotic membranes (19), POPC seemed the natural choice. To evaluate whether POPC membranes are a plausible model system for mechanistic studies, we sought to demonstrate the catalytic activity of purified hDHHC20 reconstituted into MSPE3D1 nanodiscs consisting of POPC lipids. To our knowledge, there is no prior report of any DHHC enzyme functionally reconstituted in lipid nanodiscs. Upon addition of acyl-CoA, we observed robust auto-acylation activity in samples containing hDHHC20, but no detectable activity for protein-less nanodiscs (**Fig. 2A**). Addition of the DHHC inhibitor 2-bromopalmitate (2-BP) resulted in the expected concentration-dependent inhibitory effect (**Fig. 2B**). These results demonstrate that POPC nanodiscs are valid model system to examine the mechanism of hDHHC20.

**Figure 2.**
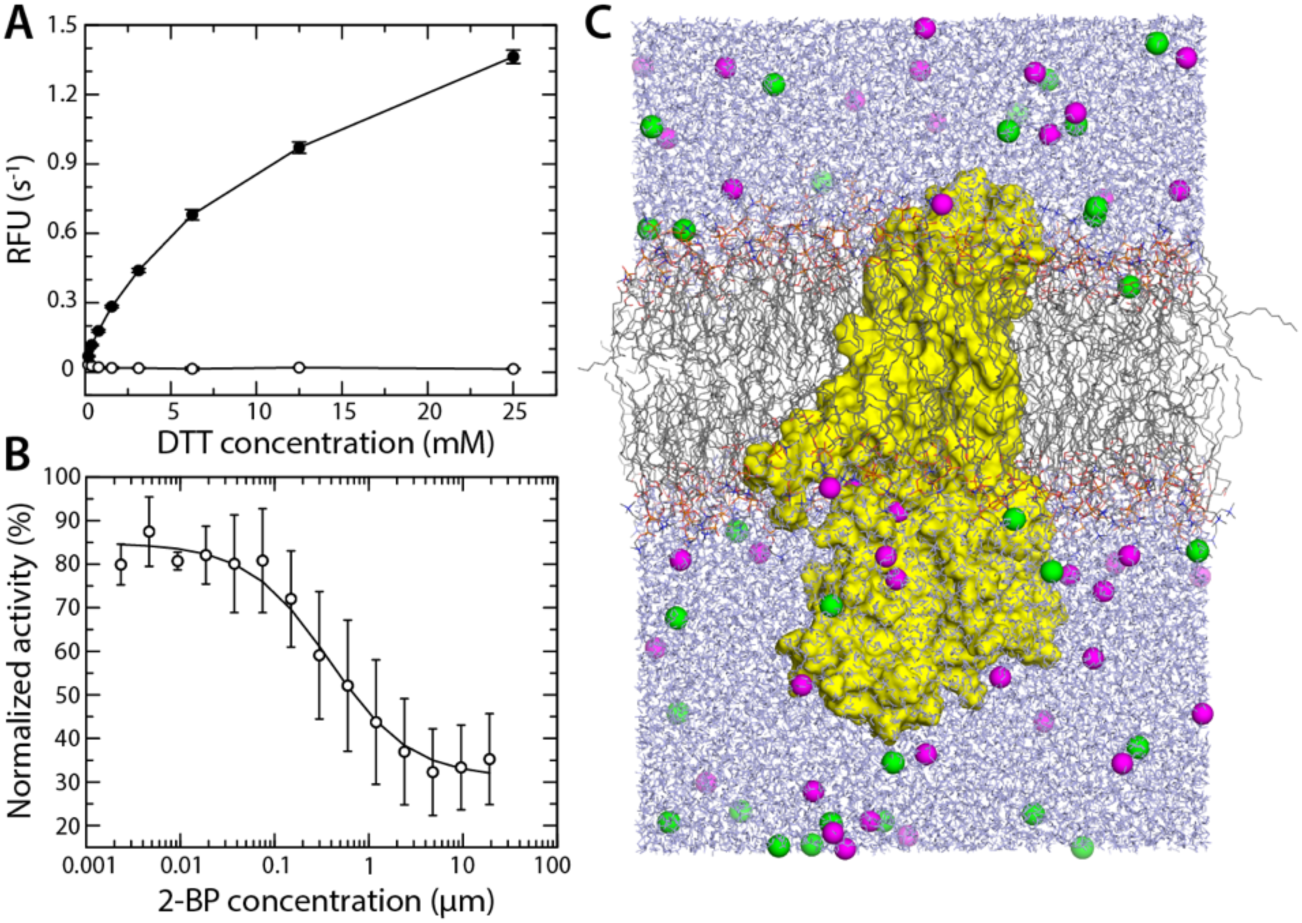
Functional reconstitution of hDHHC20 in a POPC bilayer. **(A)** Experimental assay of the auto-acylation activity of hDHHC20 in POPC nanodiscs, either in the presence of acyl-CoA (closed circles) or in its absence (open circles). Auto-acylation of hDHHC20 is spontaneous; addition of DTT reverses the reaction by breaking the thioester linkage between Cys156 and acyl-CoA, thereby regenerating the enzyme for continued catalysis. **(B)** Impairment of the auto-acylation reaction assayed in (A) by addition of 2-bromopalmitate, a competitive inhibitor of hDHHC20. **(C)** All-atom molecular dynamics simulation system, including hDHHC20 (yellow surface), a POPC bilayer (gray/blue/orange/red lines), water (blue lines) and 100 mM NaCl (green/magenta spheres). The figures shows the final snapshot of one of the two calculated trajectories, each 1-µs long.

### hDHHC20 deforms the lipid bilayer

In view of these experimental results, we prepared a simulation system for hDHHC20 in a POPC bilayer; both the protein and its environment are represented in atomic detail (**Fig. 2C**). We then carried out two independent molecular dynamics trajectories, each 1-*µ*s long. These trajectories revealed no substantial changes in the architecture of protein relative to the starting condition. For example, relative to the crystal structure, the RMS difference in the backbone of the transmembrane domain is, on average, 0.8 ± 0.1 Å, for one simulation, and 0.9 ± 0.1 Å, for the other. When all secondary-structure elements in the protein are considered, these differences are 2.2 ± 0.4 and 2.4 ± 0.6 Å.

By contrast, we observed that the structure of the lipid bilayer changed significantly in the course of both simulations. In the absence of a protein, a simulated bilayer is essentially flat, on average (**Fig. 3A**). Only small deviations are observed, due to thermal fluctuations that have not been completely averaged out. hDHHC20, however, induces a pronounced deformation in the surrounding membrane (**Fig. 3B**). This deformation is most prominent on the cytoplasmic side, where the membrane bends towards its center right in front of the catalytic site; elsewhere along the protein perimeter, the membrane is elevated instead (**Fig. 3B**). The magnitude of these perturbations is clearly significant; the displacement of the cytoplasmic ester layer in either direction is ∼5 Å, i.e. about 1/3 of the leaflet width. Accordingly, these deformations translate into a clear change in membrane width; near the catalytic site, the bilayer is approximately 8-10 Å thinner than elsewhere in the protein perimeter (**Fig. 3C**).

**Figure 3.**
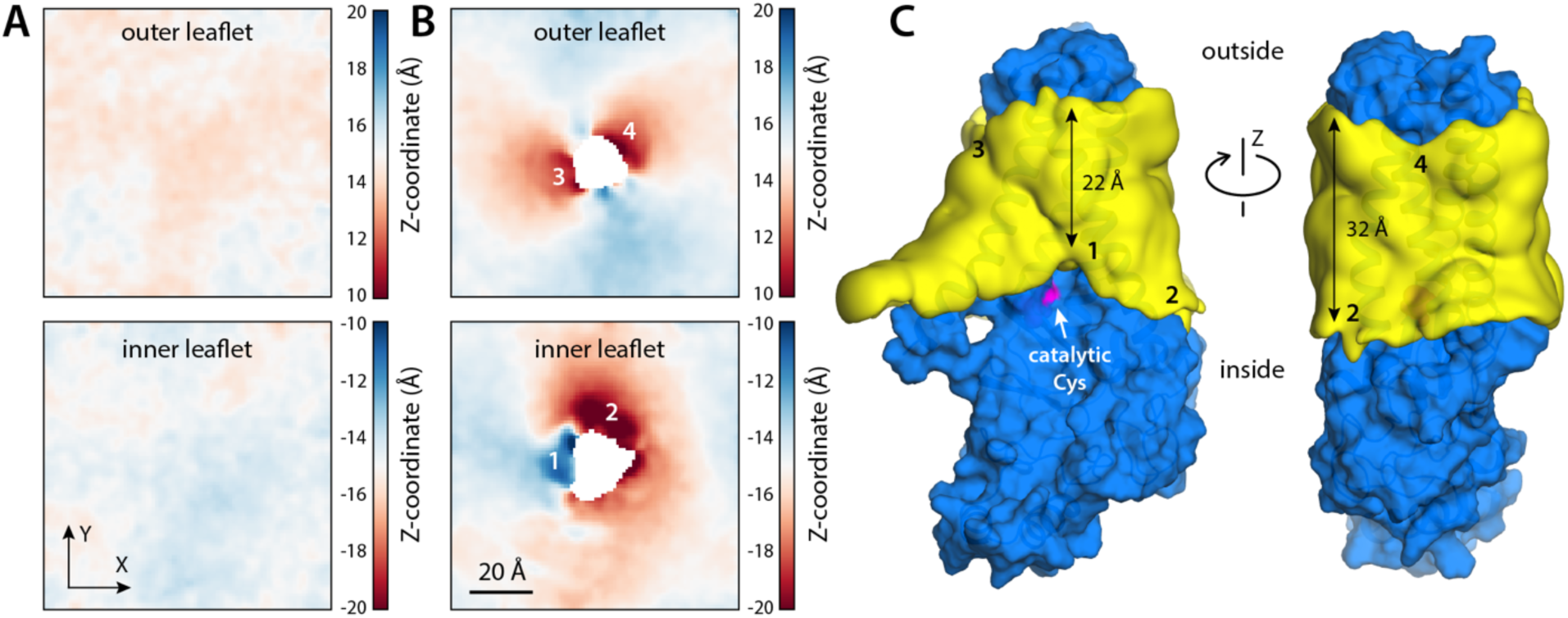
Human DHHC20 deforms the lipid bilayer to expose the catalytic cysteine. **(A)** MD simulations of a pure POPC bilayer. Having set the bilayer midplane as the XY plane (i.e. Z = 0), the plots quantify the mean Z-coordinate of the ester layer of POPC for different values of X and Y, either for the outer leaflet or the inner leaflet. The data is based on a trajectory of 330 ns. The data shows the bilayer is essentially flat, aside from minor thermal fluctuations that have not yet averaged out. **(B)** MD simulations of the same POPC bilayer with hDHHC20 embedded. The plots again quantify the mean Z-coordinate of the two ester layers in the membrane. The data is based on 2 trajectories of 1 µs each. Regions showing significant deformations are numbered. Regions, 1, 3, 4 reflect depressions, i.e. the ester layer in either leaflet bends towards the membrane center; region 2 is an elevation. **(C)** 3D density maps for the POPC alkyl-chain core (yellow surface) near the protein surface (blue), calculated from the MD trajectories. The catalytic cysteine (Cys156) is highlighted (magenta). The 4 regions where the membrane is deformed are again numbered, as in (B). The approximate minimum and maximum widths of the alkyl-chain core of the bilayer are also indicated.

The deformation of the bilayer observed in our simulations is explained by the amino-acid composition of the protein surface. Analysis of the frequency of hydrogen-bonding interactions between residues on the hDHHC20 surface and lipid molecules reveals two layers of ionized or polar side-chains that persistently associate with either the ester or phosphate layers in the POPC membrane (**Fig. 4**). The shape of this double layer of polar interactions correlates with the shape of the bilayer around the protein perimeter (**Fig. 4**), suggesting these interactions sustain the energetic cost of the observed membrane deformation.

**Figure 4.**
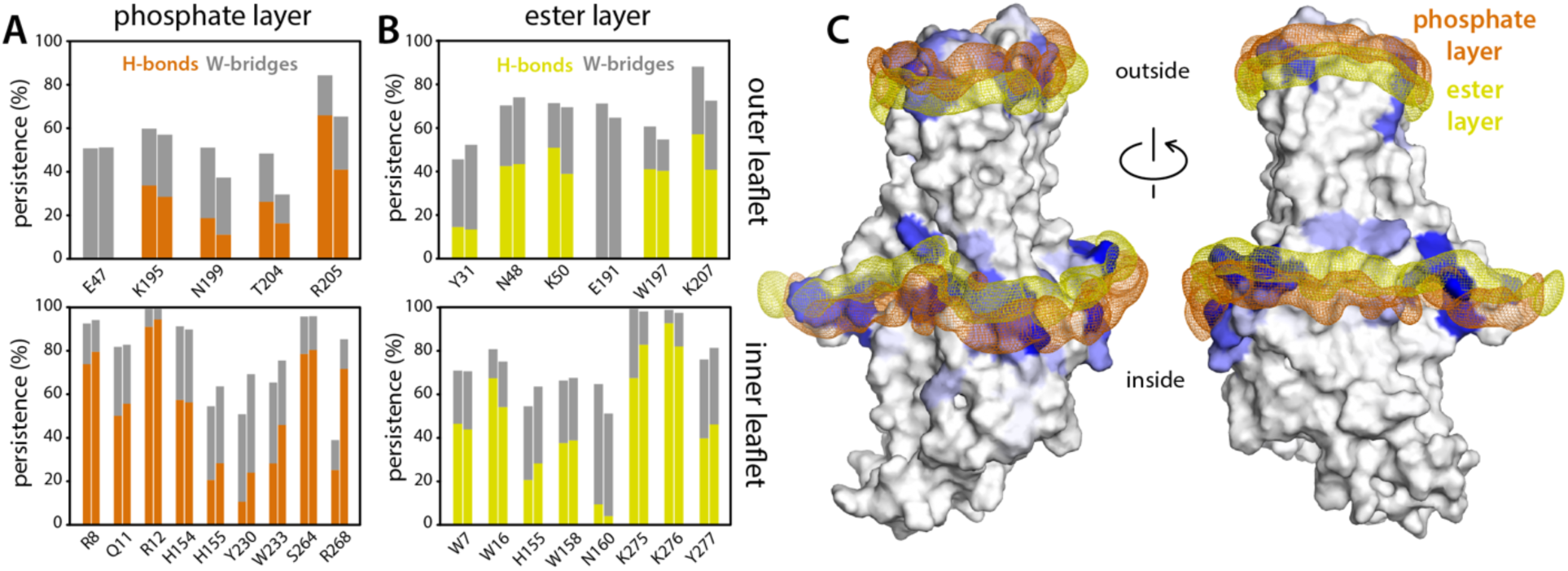
Polar interactions between hDHHC20 sidechains and surrounding lipids. **(A)** The plot quantifies the persistence of observed interactions between protein side-chains and POPC phosphate groups, i.e. the fraction of the simulated time in which the interaction was detected. The data is based on 2 independent trajectories of 1 µs each. Two types of interactions are described: direct hydrogen-bonds (H-bonds), and hydrogen-bonds bridged by 1 water molecule (W-bridge). A direct H-bond was recorded when the distance between a protein donor and a POPC acceptor is 3.2 Å or less (POPC does not contain any H-bond donors). A W-bridge was recorded when a water molecule is found to be at 3.2 Å or less of both an H-bond acceptor in POPC and either an H-bond donor or acceptor in the protein. **(B)** Same as (A), for interactions between protein side-chains and POPC ester groups. **(C)** 3D density maps for the phosphate and ester layers of POPC (orange and yellow mesh, respectively) near the protein surface, calculated from the MD trajectories. Protein residues that interact with either layer are colored in shades of blue; greater intensity denotes more persistent interactions, as quantified in (A) and (B).

### Membrane perturbation exposes catalytic Cys to water, and to reactive group in acyl-CoA

As mentioned, the structure of hDHHC20 shows Cys156 well within the hydrophobic, trans-membrane span of the protein, seemingly at odds with a putative reaction intermediate in which this side-chain is transiently ionized. The membrane perturbations revealed in our MD simulations resolve this apparent contradiction. As shown in **Fig. 5**, the localized depression of the bilayer near the catalytic site exposes Cys156 to water, to a significant degree. This result is most evident by comparison with Cys10 and Cys232, which are at the same level in the transmembrane domain but are almost entirely dehydrated. Indeed, the range of hydration numbers we observe for Cys156 overlaps signigicantly with that of Cys290, which is in the cytosolic domain and is fully exposed to solvent (**Fig. 5**).

**Figure 5.**
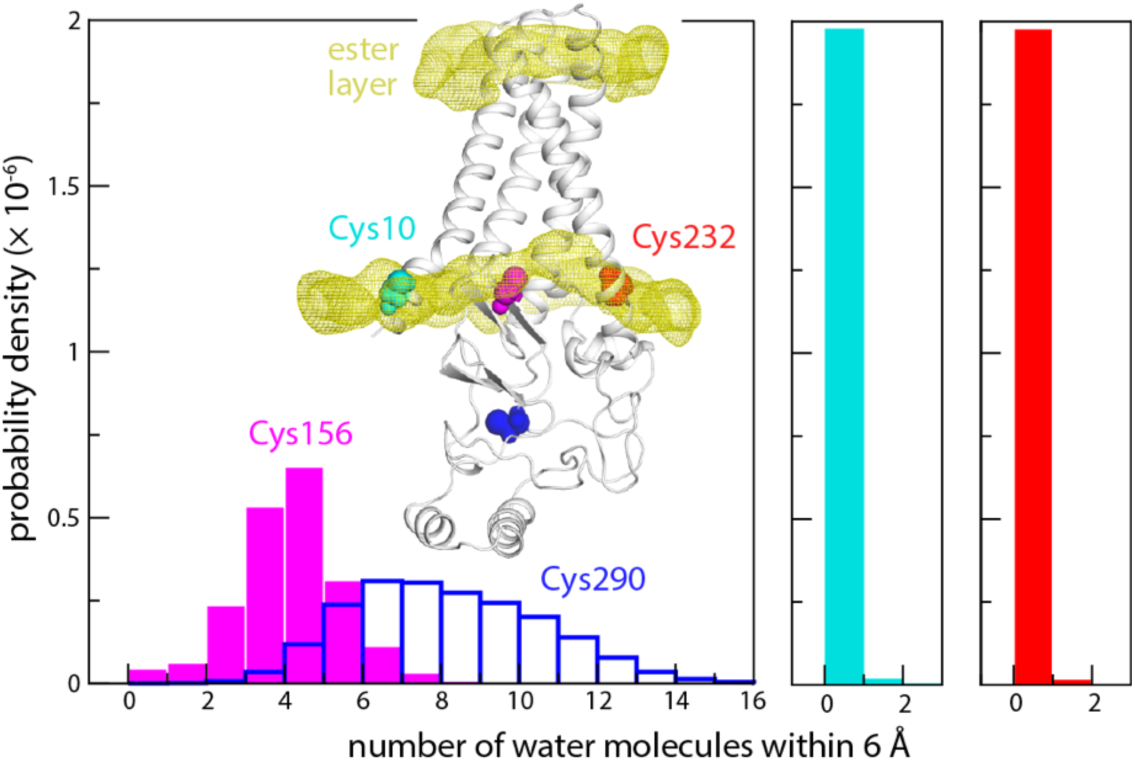
Hydration of the catalytic cysteine. The number of water molecules near Cys156 is quantified in terms of a probability distribution, derived from two MD trajectories. Analogous data is shown for a surface-exposed cysteine in the cytoplasmic domain (Cys290) and for two cysteines near the inner ester layer (Cys10, Cys232). The inset indicates the location of the 4 cysteines; calculated density maps for the two ester layers are also shown.

The depression of the membrane near the catalytic site also happens to position Cys156 at the level of the ester layer of the cytoplasmic leaflet (**Fig. 5**). We wondered whether the reactive thioester group in acyl-CoA might also tend to reside in this layer; despite the ubiquitous importance in biology of fatty acyl CoA, we could not find a previous study that addresses this question. Thus, we calculated 1-µs trajectories for molecular systems consisting of one acyl-CoA molecule in a POPC lipid bilayer, without a protein (**Fig. 6A**); we considered three acyl-chain lengths, namely C8:0 (caprylyl-CoA), C16:0 (palmitoyl-CoA) and C22:0 (behenyl-CoA). All three are substrates of hDHHC20, but as mentioned palmitoyl-CoA is the preferred substrate (10). Caprylyl-CoA and behenyl-CoA are the shortest and longest acyl-chain lengths for which hDHHC20 auto-acylation is detectable, albeit at a much smaller rate (10). Interestingly, analysis of our trajectory data shows, for all three lengths, that the reactive site in acyl-CoA is indeed aligned with the ester layer of the POPC membrane (**Fig. 6B**). The position of acyl-CoA therefore seems to be largely determined by the headgroup, rather than by the length of the hydrophobic chain. Thus, by deforming the membrane and aligning Cys156 to the ester layer, hDHHC20 appears to promote encounters between the reactive sites in the protein and substrate.

**Figure 6.**
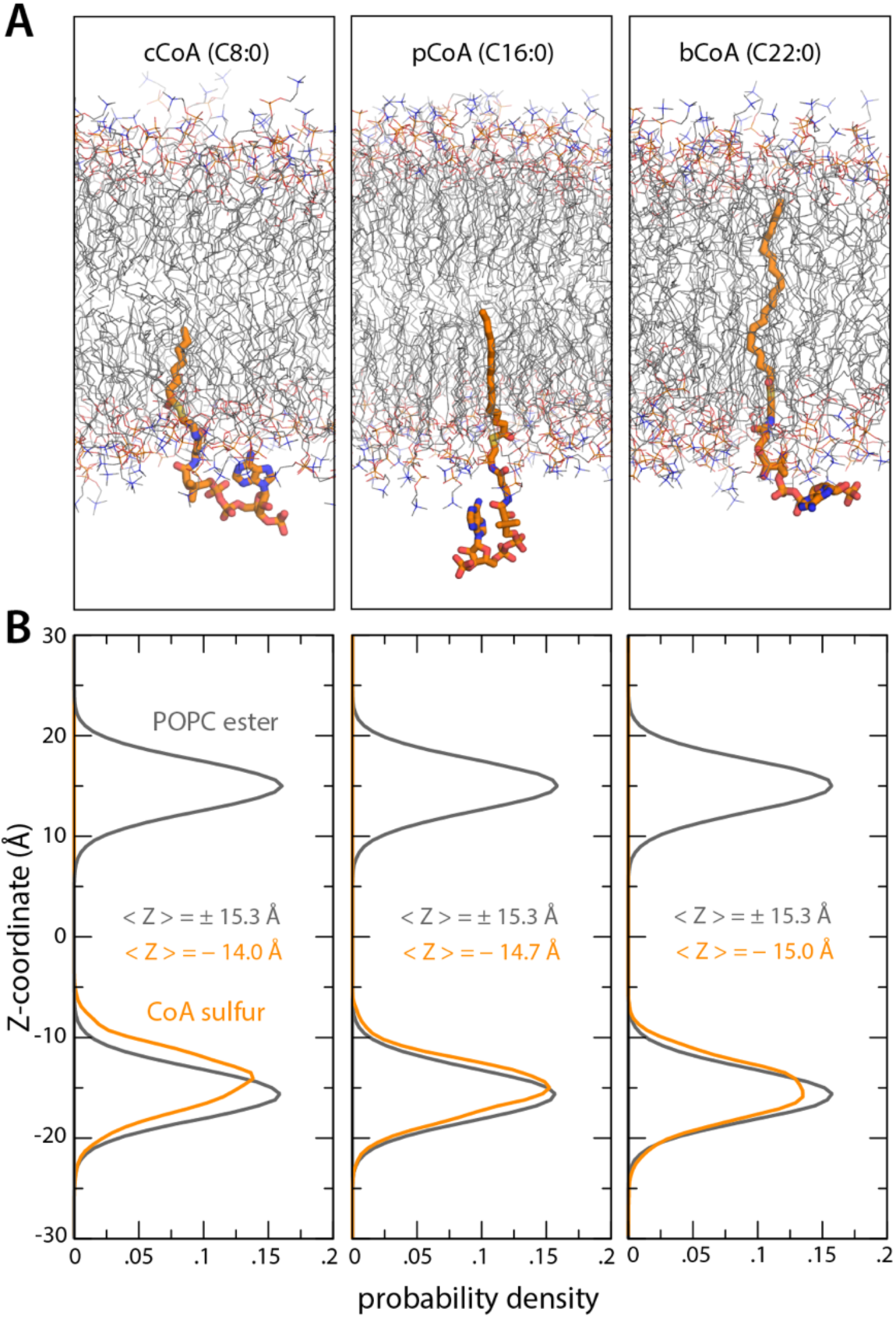
MD simulations of acyl-CoA in POPC. **(A)** Snapshots of 3 forms of acyl-CoA differing in the length of the acyl chain, from MD simulations. From left to right, these are: caprylyl-CoA (cCoA), palmitoyl-CoA (pCoA) and behenyl-CoA (bCoA). Surrounding lipid molecules are also shown (thin lines). Water, ions and hydrogens are omitted for clarity. **(B)** For each form of acyl-CoA, the plots quantify the most probable location of the sulfur atom where acyl-CoA is cleaved off (orange), relative to the membrane center and to the two ester layers in POPC. The distributions shown derive from three 1-µs MD trajectories (one for each form of acyl-CoA).

### The binding site for the acyl-CoA chain

Although no structure of DHHC bound to acyl-CoA has been reported yet, the structure of hDHHC20 inhibited by 2-bromopalmitate (2-BP) has been determined. 2-BP also reacts with Cys156 to form a covalent linkage (8). In this structure, the palmitate chain of 2-BP occupies a hydrophobic tunnel formed within the protein, on the cytoplasmic half of the transmembrane span. Because reactivity with acyl-CoA or 2-BP necessarily begins with non-covalent recognition of these ligands, we wondered whether we would observe POPC lipids bound to this site in our simulations. Indeed, analysis of the simulation data did in fact reveal that this tunnel is persistently occupied by one alkyl chain of a POPC molecule (**Fig. 7A, B**). The site is empty at the beginning of the two MD trajectories, but two different POPC lipids move to occupy the cavity early on. Interestingly, in both cases, it is the C16:0 alkyl-chain that enters the hydrophobic pocket. We next asked whether the range of configurations explored by this bound lipid during the simulations might be conducive to auto-acylation, if translated to palmitoyl-CoA. Indeed, we find that in about 5% of the configurations the distance between the reactive sites in the protein and the lipid is only 6 Å or less (**Fig. 7C**). Although this is a small population, it is worth noting that the thioester bond formed upon DHHC auto-acylation does not break spontaneously (i.e. in the absence of a thiol acceptor, in either a small-molecule or a protein); this irreversibility thus implies that auto-acylation will progress steadily even if the likelihood of reactive configurations is small. Taken together with the structure of the DHHC20-2-BP complex, these observations are strong indication that this cavity is indeed the site of recognition of the hydrophobic chain of acyl-CoA, prior to auto-acylation.

**Figure 7.**
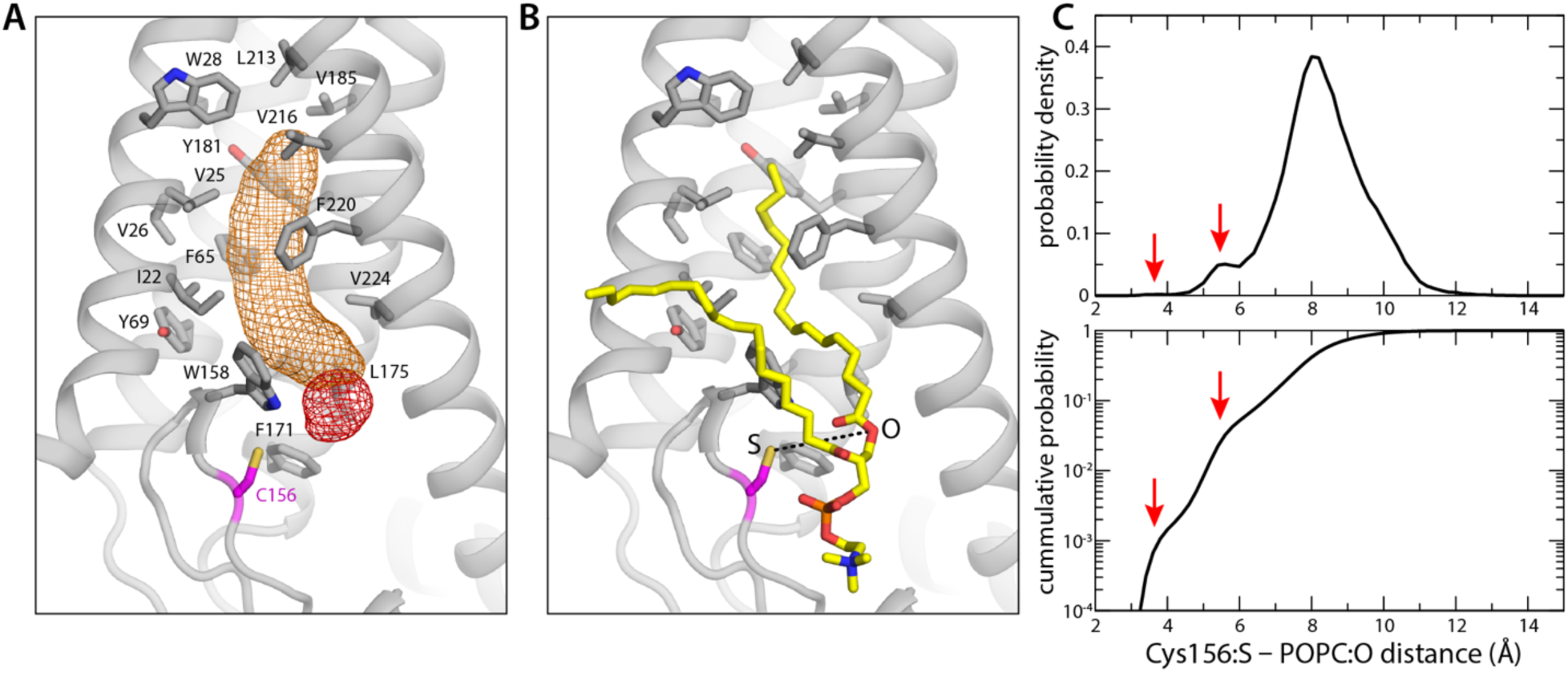
Occupancy of the putative acyl-CoA binding site by POPC. **(A)** 3D density map for the POPC alkyl-chain (orange mesh) that occupies the putative binding site for acyl-CoA throughout both of the simulated trajectories. A density map for the corresponding ester group is also shown (red mesh). The hydrophobic and/or aromatic side-chains lining the binding pocket are highlighted, as is the catalytic cysteine Cys156. The site is empty at the beginning of both simulations. Two different POPC lipids are observed to bind to the protein early on in the equilibration phase. In both cases, it is the C16:0 alkyl-chain that enters the hydrophobic pocket. **(B)** Snapshot of the bound POPC lipid, at the end of one of the two 1-µs MD trajectories (also shown in (A). In this snapshot, the distance between the sulphur atom of Cys156 and the oxygen atom in POPC that is analogous to the reactive sulfur in pCoA (dashed lines) is 8.1 Å. **(C)** Top panel: probability distribution for the S-O distance described in (B), calculated from all trajectory data; lower panel: integral of this probability distribution. Red arrows indicate low-population states that would likely foster the formation of a thioester bond, if acyl-CoA was bound as POPC.

## Discussion and Conclusions

Despite the physiological importance of protein palmitoylation, little is known about the molecular mechanism of the enzymes that catalyze this post-translational modification, known as DHHC acyl-transferases. In a breakthrough, the atomic structure of human DHHC20 was recently determined, both for an apo state as well as in complex with the inhibitor 2-BP (8). These structures provided the necessary framework to generate atomically-detailed mechanistic hypotheses, but also opened up a range of new questions. Particularly pertinent among them are the nature of the interactions between DHHC enzymes and the surrounding membrane, and how the membrane, in turn, defines the chemical enviroment around the active site. This study builds upon that discovery, and seeks to begin to provide specific insights into these questions.

More specifically, we first aimed to establish an experimental in vitro system to evaluate the activity of the enzyme in the context of a lipid membrane, rather than in detergent, which had been the condition of previous functional assays (8). This development was essential since S-acylation requires that DHHC proteins first recognize acyl-CoA molecules residing in the membrane. The fatty acyl group then becomes covalently linked to a membrane-proximal cysteine, and is ultimately transferred to a substrate protein. The membrane thus appears to be an integral element of this process. A nanodisc system such as that developed here will enable us to examine a range of mechanistic aspects, such as the molecular basis for the varying substrate specificities of different DHHC enzymes, or how the lipid composition of the membrane defines catalytic rates. Another area of interest is of course the interaction of the enzyme with potential drugs; indeed, two human DHHC have been proposed as targets for novel therapeutic interventions against cancer (20).

Our second aim was to begin to understand the interplay between DHHC enzymes and the membrane, in atomic detail. To this end, we resorted to molecular dynamics simulations, which had not been previously employed to study this class of enzymes. The simulations enabled us to clarify a puzzling observation in the crystal structure of hDHHC20, namely that the catalytic cysteine (Cys156) is well within the hydrophobic transmembrane span of the protein. Shielded from water, the transient ionization of Cys156, which must precede the formation of the thioester bond with the chain of acyl-CoA, would be highly disfavored. The answer to this puzzle is that hDHHC20 induces a pronounced depression of the membrane near the catalytic site, precisely to expose Cys156 to water. Interestingly, our simulations show that this perturbation of the membrane also aligns Cys156 with the ester layer, which is precisely where the cleavage site in acyl-CoA resides. These insights underscore the notion that the interplay between membrane and protein is an integral element of the mechanism of this class of enzymes. These results also provide a solid foundation for future simulation studies; for example, a direct examination of the process of recognition of acyl-CoA ought to reveal important clues into the determinants of the chain-length specificity of hDHHC20.

The ligands of many membrane-integral enzymes and transporters are hydrophobic (or at least amphipathic), implying these ligands are most likely recognized (and released) laterally, from (and to) the membrane instead of through the aqueous phase. In particular membrane-integral enzymes with lipid-like ligands include many transferases(20-24), as well as some translocases(25), synthases(26), polymerases (27) and proteases(28). Little is known about how most of these proteins interact with the membrane, but the increasing availability of structural data will surely help advance our understanding. For example, in the structure of the isoprenylcysteine carboxyl methyl-transferase (ICMT), the arrangement two of the transmembrane helices appears to indicate a depression in the lipid membrane near the active site, to help accommodate the substrates (20). Similar membrane perturbations have been noted for the rhomboid protease GlpG based on pioneering MD simulations (29, 30), later corroborated by EPR spectroscopy (31). Taken together with our observations for DHHC20, the paradigm that seems to be emerging for these types of membrane proteins is that they have evolved highly specific ways to perturb the morphology of the surrounding lipid bilayer, precisely to facilitate the recognition of substances and substrates that reside therein.

## Author contributions

R.S. performed the computational work and analyzed the data; J.S. performed the experimental work and analyzed the data; A.B. designed and supervised the experimental work; J.D.F-G. designed and supervised the experimental work and data analysis. All authors contributed to write the manuscript and prepare figures.

## Acknowledgements

This work was funded by the Divisions of Intramural Research of the National Heart, Lung and Blood Institute (R.S. and J.D.F-G.) and the National Institute of Child Health and Human Development (J.S. and A.B), National Institutes of Health (NIH), USA. Computational resources were in part provided by the NIH HPC facility Biowulf.

